# Exercise training remodels inter-organ endocrine networks

**DOI:** 10.1101/2025.04.29.651323

**Authors:** Cheehoon Ahn, Andrea L. Hevener, Laurie J. Goodyear, Sue C. Bodine, Karyn A. Esser, Marcus M. Seldin, Lauren M. Sparks

## Abstract

Exercise induces organism-wide molecular adaptations, partly mediated by humoral factors released in response to acute and chronic physical activity. However, the extent and specificity of endocrine effects from training-induced secreted factors remain unclear. Here, we applied systems genetics approaches to quantify inter-organ endocrine networks using multi-tissue transcriptomics and proteomics data collected from endurance-trained rats in The Molecular Transducers of Physical Activity Consortium (MoTrPAC). Eight weeks of endurance training significantly altered both the magnitude and specificity of endocrine effects across multiple origin-target tissue pairs. Subcutaneous white adipose tissue emerged as a key endocrine regulator impacted by training, while extracellular matrix-derived factors were identified as globally regulated secretory features in trained vs sedentary animals. Notably, secretory Wnt signaling factors were identified as key mediators of exercise-induced endocrine adaptations in multiple tissues. Our systems genetics framework provides an unprecedented atlas of inter-organ communication significantly remodeled by endurance exercise, serving as a valuable resource for novel exerkine discovery.

## INTRODUCTION

Exercise enhances cardiometabolic, immune, and neurological health (1). These health benefits are accompanied by multi-organ molecular adaptations, partially mediated by humoral exerkines—factors released from various tissues following physical activity (2, 3). In addition, daily physical activity or daily exercise may induce chronic alterations in inter-organ communication that extend beyond the acute exerkine response (4). Recent advancements in secretome profiling have enabled identification of the cellular and tissue origins of acutely induced exerkines (5), greatly expanding our understanding of exercise-stimulated humoral signaling. Furthermore, inter-tissue correlations of exercise-responsive metabolomes show a tissue-pair co-regulation (6). However, the full extent of exerkine-mediated endocrine effects (i.e., magnitude and target tissue specificity) and the broader exercise-stimulated remodeling of the inter-organ endocrine network remain inadequately understood. Methodological limitations have hampered progress in addressing these research questions.

To address the gap in research capability for exerkine discovery, we employed Quantitative Endocrine Network Interaction Estimation (QENIE) (7, 8), a systems genetics approach that leverages gene-to-gene correlations across tissue pairs, which has been successfully implemented to identify novel tissue crosstalk factors (7, 9, 10). QENIE assigns a secretome score (S_sec_) to each known secreted feature within a given origin-target tissue pair, providing a quantitative estimate of its endocrine impact. Additionally, we recently developed Gene-Derived Correlations Across Tissues (GD-CAT) (11), a bioinformatics pipeline that performs gene set testing on target tissue genes that correlate with a given gene in an origin tissue. This approach allows us to infer biological pathways in the target tissue that may be modulated by the endocrine action of the origin tissue gene.

The Molecular Transducers of Physical Activity Consortium (MoTrPAC) recently reported temporal multi-omic adaptations in response to eight weeks of endurance training intervention across multiple solid tissues in male and female rats—representing the most comprehensive preclinical exercise training study to date (12, 13). Leveraging multi-organ transcriptomics and proteomics data from the MoTrPAC rat endurance training study, combined with the robust analytical power of QENIE and GD-CAT, we aimed to quantify endurance training-induced inter-organ endocrine networks and identify novel secreted factors mediating tissue crosstalk. We hypothesized that QENIE and GD-CAT would recapitulate well-established endocrine interactions known to be impacted by exercise (e.g., leptin signaling from adipose tissue to the hypothalamus). Additionally, we predicted that trained rats would exhibit distinct inter-organ endocrine networks compared with sedentary control rats, reflecting systemic adaptations to daily treadmill running.

## RESULTS

### Application of QENIE and GD-CAT

We first evaluated whether inter-tissue transcriptomic correlations in MoTrPAC rats were primarily reflected shared biological processes across tissues, rather than direct tissue-to-tissue regulatory interactions. We performed Weighted Gene Co-expression Network Analysis (WGCNA) on 16 tissues (ovary and testis were excluded due to limited sample sizes) using all available samples from an exercise timecourse intervention from 1-8 weeks (nL=L10 for each training group; 1-week, TR1W; 2-week, TR2W; 4-week, TR4W; 8-week, TR8W; 8-week untrained control, CON; total 50 samples per tissue), resulting in 203 unique tissue-specific gene modules (see ‘WGCNA’ in Methods). Correlations between module eigengenes revealed several significant inter-tissue relationships (Figure S1A). However, overrepresentation analysis showed that even highly correlated modules were often enriched for distinct biological pathways (Figure S1A). To determine whether functional similarity explained module correlations, we calculated Jaccard indices based on overlapping significant Gene Ontology (GO) terms between modules. The resulting similarity pattern (Supplementary Figure 1B) did not mirror the eigengene correlation structure, suggesting that inter-tissue transcriptomic correlations were not driven by common pathway enrichment. Moreover, genes encoding secretory proteins—defined using the UniProt secretome database (14)—were broadly distributed across modules and tissues, with no clear clustering of secretory content in a specific module or tissue (Supplementary Figure 1A). These findings suggest that the observed inter-tissue transcriptomic correlations are unlikely to be explained by shared biological pathways across tissues alone.

The workflow and implementation of QENIE (7) are outlined in Figure 1A. Biweight midcorrelation (15) was performed between the gene matrix from the origin tissue and the target tissue. From the resulting p-value matrix, secretory protein-coding transcripts in the origin tissue were selected (see QENIE in Methods), yielding a varied number of secretory protein-coding transcripts per origin tissue (Figure 1B). The secretome score (S_sec_) for each origin transcript was then calculated as the sum of -log(p-value) for its correlations with all target tissue transcripts. The key assumption underlying QENIE is that if the transcript abundance of a secreted protein-coding gene in the origin tissue correlates with transcripts in the target tissue, then the effector gene may act as a mediator of inter-tissue crosstalk. Therefore, S_sec_ quantifies the potential endocrine effect of a given transcript on its target tissue. QENIE was applied across all possible gene-to-gene correlations of origin-target tissue pairs. S_sec_ values were also calculated from protein-to-protein correlations in seven of the 16 available tissues, producing 1,525 unique S_sec_ datasets, including matched origin-target pairs (i.e., auto/paracrine actions) (Figure 1A). The number of filtered secretory features averaged 812 ± 78 for transcripts and 475 ± 88 for proteins across origin tissues (Figure 1B). Notably, the extent of differential regulation in response to 8 weeks of endurance training (FDR < 0.05; available here: https://motrpac-data.org/search) of secretory features varied widely across tissues. For example, 44% (351 out of 793) of secretory protein-coding transcripts in the adrenal gland were differentially regulated by endurance training, whereas only 0.001% of secretory protein-coding transcripts in the hypothalamus exhibited training-induced changes (Figure 1B). At the protein level, 90 secretory proteins in subcutaneous white adipose tissue (scWAT) were differentially regulated by exercise, while only 1 secretory protein in the cerebral cortex was altered following exercise intervention compared with sedentary control (Figure 1B). The raw abundance of secretory features was also partly tissue-dependent (Supplementary Figure 2A, B).

**Figure 1.**
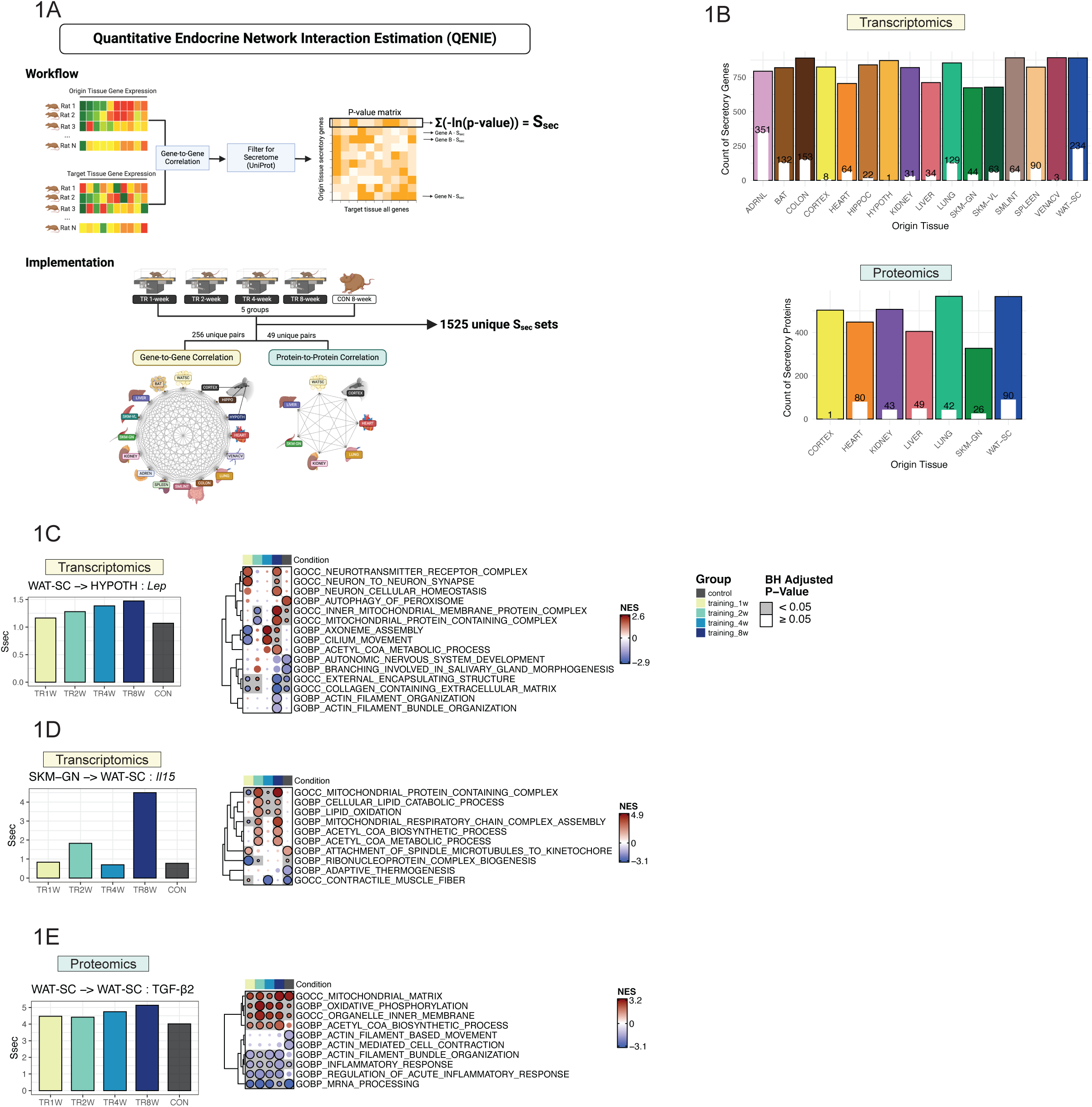
Workflow of QENIE and validation of physiological relevance of S_sec_. A) Schematic representation of the QENIE workflow and its implementation. B) Number of secretory transcripts and proteins identified in each origin tissue after secretome filtering. White bars and numbers within the colored bars indicate the count of secretory transcripts or proteins that were significantly different between CON and TR8W. C) S_sec_ of Lep (WAT-SC-to-HYPOTH) across groups. Fgsea results of HYPOTH transcripts correlating with WAT-SC-*Lep* across groups are shown. D) S_sec_ of Il15 (SKM-GN-to-WAT-SC) across groups. Fgsea results of WAT-SC transcripts correlating with SKM-GN-*Il15* across groups are shown. E) S_sec_ of TGF-β2 (WAT-SC-to-WAT-SC) across groups. Fgsea results of WAT-SC proteins correlating with WAT-SC-TGF-β2 across groups are shown. QENIE, Quantitative Endocrine Network Interaction Estimation; S_sec_, Secretome score; ADRNL, Adrenal gland; BAT, brown adipose tissue; COLON, colon; CORTEX, cerebral cortex; HEART, heart; HIPPOC, hippocampus; HYPOTH, hypothalamus; KIDNEY, kidney; LIVER, liver; LUNG, lung; PLASMA, plasma; SKM-GN, gastrocnemius (skeletal muscle); SKM-VL, vastus lateralis (skeletal muscle); SMLINT, small intestine; SPLEEN, spleen; VENACV, vena cava; WAT-SC, subcutaneous white adipose tissue. CON, Control; TR, Training; GOBP, Gene Ontology Biological Process; GOCC, Gene Ontology Cellular Component; GOMF, Gene Ontology Molecular Function; NES, Normalized Enrichment Score; BH, Benjamini-Hoechberg.

To assess the physiological relevance of S_sec_, we examined well-established tissue crosstalk factors that are modulated by exercise: leptin (*Lep*), IL15 (*Il15*), and TGF-β2 (3, 16). Leptin, a hormone predominantly secreted by scWAT (Supplementary Figure 2A), plays a critical role in regulating energy homeostasis, with exercise training known to enhance hypothalamic sensitivity to leptin (17, 18). S_sec_ of *Lep* (scWAT-to-hypothalamus) increased progressively throughout training, with the most pronounced difference observed between TR8W and CON (Figure 1C). To infer the potential biological effects of scWAT-derived leptin in the hypothalamus, Gene-Derived Correlation Across Tissue (GD-CAT) was performed (11). While scWAT-*Lep* was associated with downregulated extracellular matrix (ECM) pathways in the hypothalamus across both groups, it was uniquely correlated with upregulated neuronal synapse and homeostasis, and neurotransmitter receptor complex in the hypothalamus of TR8W, supporting the notion that endurance training may contribute to enhanced synaptic plasticity of the hypothalamus (adjusted p<0.05, Figure 1C). IL15 is a well-characterized exercise-inducible myokine that promotes triglyceride breakdown (i.e., lipolysis) and fatty acid oxidation in adipose tissue (19, 20). Compared with CON, S_sec_ of *Il15* (gastrocnemius-to-scWAT) was over four-fold higher in TR8W (Figure 1D). Additionally, genes in scWAT that correlated with gastrocnemius-*Il15* in TR8W exhibited a significant upregulation of lipid catabolism and oxidation, an effect absent in CON, aligning with the known role of IL15 in regulating adipose tissue energy metabolism (adjusted p<0.05, Figure 1D). TGF-β2, an adipokine reported to be induced by exercise, improves adipose tissue metabolism and reduces inflammation (21). S_sec_ of TGF-β2 protein (scWAT-to-scWAT) increased with training (Figure 1E), and GD-CAT analysis revealed that scWAT-TGF-β2 was generally associated with upregulated mitochondrial metabolism and reduced inflammatory signaling in scWAT regardless of training state (Figure 1E). However, some pathways were differentially enriched between CON vs. TR8W, including the upregulation of acetyl-CoA biosynthesis and the downregulation of acute inflammatory response in TR8W (Figure 1E). Collectively, our *in silico* observations from QENIE and GD-CAT analyses support the known endocrine roles of selected exercise-stimulated features and expand our understanding of the role of endurance training in remodeling the endocrine network.

### Endurance training alters S_sec_ ranks and distribution

To assess the overall endocrine contribution of each origin tissue, we ranked tissues based on median S_sec_ from gene-to-gene correlations. Notably, vena cava exhibited the highest median S_sec_ at TR1W and TR2W, but by TR4W and TR8W, scWAT emerged as the top-ranked origin tissue (Supplementary Figure 3A). Mean S_sec_ was significantly different in 187 out of 256 gene-to-gene origin-target pairs and 32 out of 49 protein-to-protein pairs between TR8W and CON (adjusted p<0.05, Supplementary Figure 3B, C). To determine the impact of endurance training on S_sec_ ranking and distribution of individual secretory features, we performed Wilcoxon signed-rank tests combined with rank sum effect sizes. In gene-to-gene correlations, 186 origin-target pairs exhibited significantly different S_sec_ ranks between TR8W and CON (adjusted p<0.05, Supplementary Figure 4A), with scWAT-to-vastus lateralis showing the highest significance and effect size. These findings indicate that adipose-to-skeletal muscle crosstalk was most differentially regulated by training (Figure 2A). Several scWAT-to-tissue connections, such as scWAT-to-brown adipose tissue (BAT) and scWAT-to-hypothalamus, were also among the top-ranked interactions, suggesting that scWAT is a primary endocrine tissue highly influenced by endurance training (Figure 2A). S_sec_ distributions were markedly skewed toward higher S_sec_ in scWAT transcripts in TR8W vs. CON, further supporting this finding (Supplementary Figure 4B). In contrast, small intestine-to-vastus lateralis, small intestine-to-colon, and vena cava-to-hippocampus exhibited skewed distributions toward lower S_sec_ in TR8W, suggesting a diminished endocrine link after training (Supplementary Figure 4C, D). Among matched origin-target pairs, small intestine and hypothalamus were most differentially regulated, with both exhibiting a shift towards higher S_sec_ in TR8W vs. CON (Supplementary Figure 4F).

**Figure 2.**
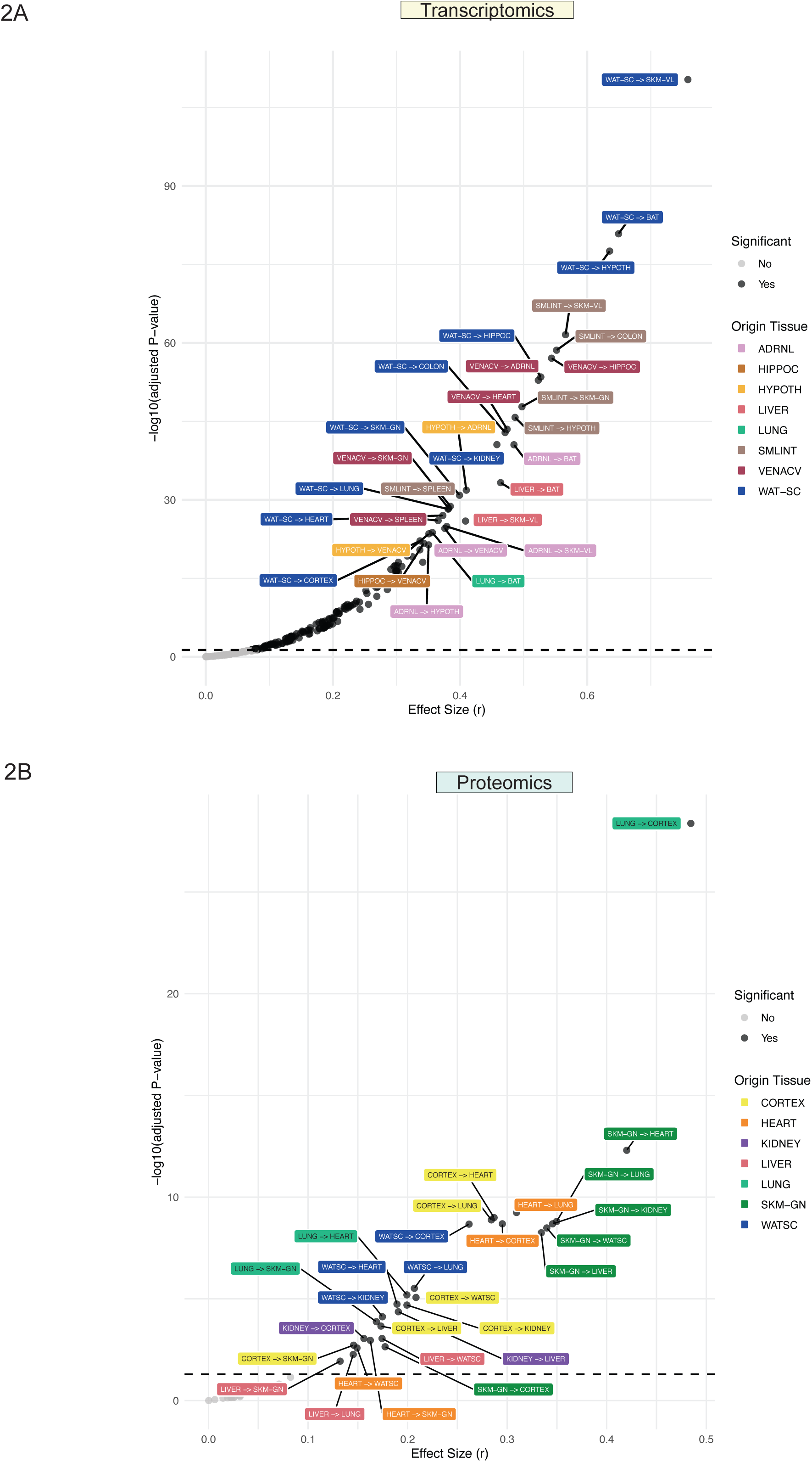
S_sec_ rank difference in CON vs. TR8W. A) Volcano plot showing results of Wilcoxon signed-rank test comparing gene-to-gene S_sec_ rank differences between CON and TR8W. B) Volcano plot showing results of Wilcoxon signed-rank test comparing protein-to-protein S_sec_ rank differences between CON and TR8W. Significant (adjusted p<0.05) origin-target pairs are highlighted in black. Color-coded padded boxes represent different origin tissues. Matched origin-target pairs (e.g., CORTEX→CORTEX) are excluded from A and B but were included in the statistical testing.

For protein-to-protein correlations, 31 out of 49 origin-target pairs showed significantly different S_sec_ ranks between TR8W and CON, with lung-to-cerebral cortex and gastrocnemius-to-heart being the most differentially regulated connections (Figure 2B). scWAT and gastrocnemius were also among the most differentially regulated matched origin-target pairs, suggestive of differential autocrine actions by endurance training (Supplementary Figure 4G, H).

### Global and tissue-specific S_sec_ changes by endurance training

Given the marked differences observed in gene-to-gene and protein-to-protein correlation networks between TR8W and CON, we examined the most differentially ranked secretory protein-coding transcripts that were globally expressed across all 16 tissues. Among 524 globally expressed transcripts, the top 20 with the greatest S_sec_ rank differences between TR8W and CON included *Adamts3, Adamtsl4, Cd9, Col12a1, Adamts4, Ctss, Endou, Fstl3, Tsku, Calr, Reln, Klkb1, Egfl7, Fam3a, Itgbl1, Mbp, Rarres2, Gpi, Lgals3,* and *Loxl2* (Supplementary Figure 5A). Notably, many of these transcripts exhibited tissue-specific differential regulation in response to training (FDR<0.05, Supplementary Figure 5B). Several of these transcripts were involved in extracellular matrix (ECM) construction and remodeling, including *Adamts3, Adamtsl4, Col12a1, Endou,* and *Loxl2*, all of which exhibited significant differences in average S_sec_ between TR8W and CON (all p<0.05, Figure 3A).

**Figure 3.**
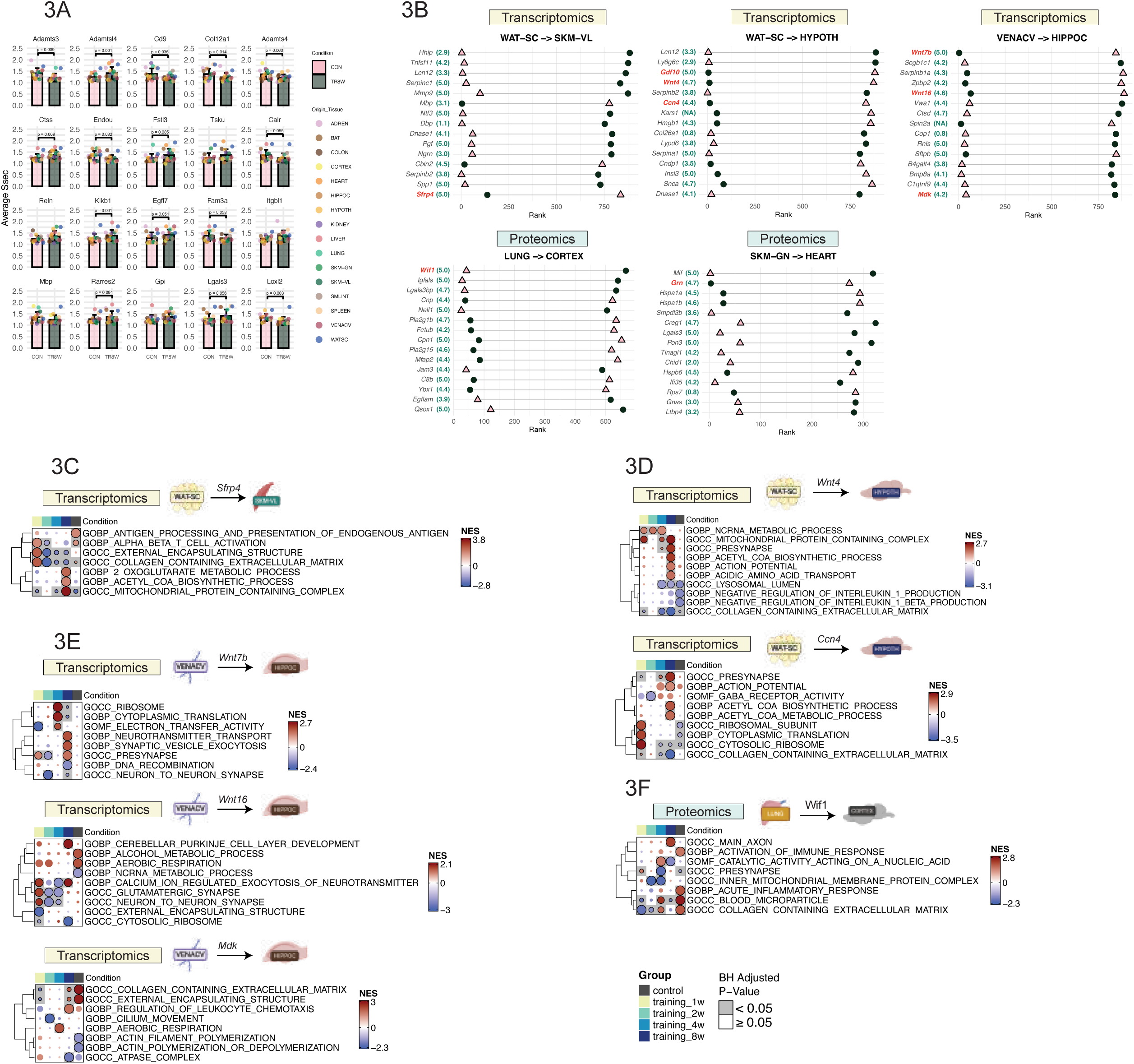
Top secretory genes with the largest S_sec_ rank difference in CON vs. TR8W. A) Bar plots displaying the results of a paired t-test comparing S_sec_ values between CON and TR8W for the top 20 global transcripts exhibiting the largest S_sec_ rank differences between the two groups. B) Top 15 transcripts exhibiting the largest S_sec_ rank difference between the CON and TR8W for WAT-SC-to-SKM-VL, WAT-SC-to-HYPOTH, VENACV-to-HIPPOC, LUNG-to-CORTEX, and SKM-GN-to-HEART. Extracellular score (range: 0-5) obtained from COMPARTMENTS is shown in green numeric next to each feature. Pink triangles represent S_sec_ ranks in CON, and dark green circles represent S_sec_ ranks in TR8W. Features highlighted in subsequent figures are colored in red. C) Fgsea results of SKM-VL transcripts correlating with WAT-SC-*Sfrp4* across groups. D) Fgsea results of HYPOTH transcripts correlating with WAT-SC-*Wnt4* and WAT-SC-*Ccn4* across groups. E) Fgsea results of HIPPOC transcripts correlating with VENACV-*Wnt7b*, VENACV-*Wnt16*, and VENACV-*Mdk* across groups. FFgsea results of CORTEX proteins correlating with LUNG-Wif1 across groups.

We then examined secretory protein-coding features that displayed substantial S_sec_ rank differences at the individual tissue level, focusing on top-ranked features with the largest shifts between TR8W and CON across key origin-target tissue pairs. Additionally, we mapped an “extracellular score” (range: 0-5, with higher scores indicating higher secretion probability) for each feature using COMPARTMENTS (22), a subcellular localization database (see ‘Extracellular score’ in Methods) (Figure 3B). In the scWAT-to-hypothalamus connection, *Gdf10* (growth differentiation factor 10, extracellular score: 5) exhibited a dramatic difference in S_sec_ rank, rising from 880^th^ in CON to 6^th^ in TR8W (Figure 3B). GD-CAT analysis revealed that scWAT-*Gdf10* was associated with reduced ECM collagen abundance and cytoskeletal remodeling in the hypothalamus of TR8W, effects absent in CON (adjusted p<0.05, Supplementary Figure 5C). In protein-to-protein correlations, Granulin (Grn, extracellular score: 4.7), a muscle growth regulator (23), increased in S_sec_ rank from 274^th^ in CON to 3^rd^ in TR8W in the gastrocnemius-to-heart connection (Figure 3B). GD-CAT linked gastrocnemius-derived Grn to enhanced amino acid metabolism and reduced RNA splicing in the heart of TR8W, an effect not observed in gastrocnemius-to-heart of CON (adjusted p<0.05, Supplementary Figure 5D).

Among scWAT-to-vastus lateralis, scWAT-to-hypothalamus, vena cava-to-hippocampus, and lung-to-cortex, Wnt signaling factors (*Sfrp4, Wnt4, Ccn4, Wnt7b, Wnt16, Mdk, Wif1*), exhibited large S_sec_ rank shifts in TR8W vs CON (Figure 3B). Wnt signaling, an essential development pathway mediated by secreted Wnt family ligands (24), is known to exert tissue-specific regulatory effects (25, 26). *Sfrp4* (Secreted Frizzled-Related Protein 4), a Wnt signaling modulator, showed a positive shift in S_sec_ rank from CON to TR8W (Figure 3B). GD-CAT linked scWAT-*Sfrp4* to increased mitochondrial metabolism in the vastus lateralis of TR8W, while it was associated with upregulated immune cell activation pathways in CON (adjusted p<0.05, Figure 3C). Interestingly, although Sfrp4 expression in scWAT was significantly downregulated with training (FDRL=L9.6e-8), its Ssec rank increased, suggesting that lower Sfrp4 expression may enhance its relative endocrine influence on vastus lateralis during training. Our observation of upregulated mitochondrial metabolism in the vastus lateralis with reduced scWAT-*Sfrp4* aligns with previous findings that Sfrp4 treatment of myotubes blunts mitochondrial respiration (27).

In the scWAT-to-hypothalamus connection, both *Wnt4* (extracellular score: 4.7) and *Ccn4* (extracellular score: 4.4) exhibited higher S_sec_ ranks in TR8W than CON, correlating with enhanced mitochondrial metabolism, synaptic activity, and action potential regulation in the hypothalamus (adjusted p<0.05, Figure 3D). In the vena cava-to-hippocampus connection, *Wnt7b* (extracellular score: 5) and *Wnt16* (extracellular score: 4.6) were among the most highly ranked secretory transcripts in TR8W compared with CON (Figure 3B). *Wnt7b* was linked to enhanced neurotransmitter secretion, while *Wnt16* was associated with upregulated Purkinje cell development and neurotransmitter exocytosis in the hippocampus, effects absent in CON (adjusted p<0.05, Figure 3E). In contrast, *Mdk* (Midkine, extracellular score: 4.2), a negative Wnt regulator, exhibited a large negative S_sec_ shift in TR8W, suggesting a weakened endocrine role (Figure 3B). Vena cava-derived *Mdk* was associated with ECM accumulation in both TR8W and CON, but only correlated with suppressed actin polymerization terms in CON (adjusted p<0.05, Figure 3E), suggesting that training may have attenuated the inhibitory effects of *Mdk* on cytoskeleton remodeling. At the protein level, Wif1 (Wnt Inhibitory Factor 1, extracellular score: 5) exhibited the largest negative S_sec_ rank shift in lung-to-cerebral cortex, with a marked reduction in TR8W (Figure 3B). In the cerebral cortex, lung-derived Wif1 was linked to axonal growth in TR8W, while in CON, it was associated with inflammation and ECM accumulation (adjusted p<0.05, Figure 3F). These findings suggest that exercise training may attenuate lung-derived Wif1’s endocrine role, potentially reducing neuroinflammatory pathways in the brain.

Because exercise training has been reported to upregulate Wnt signaling in the brain (28, 29), we investigated whether the observed training-induced inter-organ communication was a secondary effect of autocrine or paracrine Wnt signaling within the brain. Interestingly, Wnt connections within the hypothalamus and hippocampus (autocrine/paracrine signaling) often showed biological pathway associations that differed markedly—and in some cases, were opposite—from those inferred through distant endocrine connections. For instance, hypothalamic-derived *Wnt4* and *Ccn4* were negatively associated with action potential and synaptic activity in the hypothalamus (adjusted p<0.05, Supplementary Figure 6A). Notably, *Ccn4* exhibited a dramatic drop in S_sec_ rank, falling from 8^th^ in CON to 615^th^ in TR8W, suggesting that its role in intra-hypothalamic signaling may have diminished following training. Furthermore, hippocampus-derived *Ccn4* showed no significant pathway enrichment in the hippocampus-to-hypothalamus connection (Supplementary Figure 6A). Similarly, while hippocampus-derived *Wnt7b* was associated with upregulated synapse signaling in TR8W, this effect was already significant in CON (Supplementary Figure 6B), indicating that local *Wnt7b* may not be part of the hippocampal synaptic remodeling differentially induced by exercise training. These findings suggest that endocrine Wnt signaling across distant tissues may not be a secondary consequence of autocrine or paracrine Wnt signaling within the brain.

### Secretory Wnt factors as central mediators of training-induced endocrine remodeling

Given the strong involvement of Wnt factors in training-induced endocrine remodeling, we systematically assessed positive, negative, and dual Wnt regulators, comparing their average S_sec_ ranks between TR8W and CON (see “Curation of secretory Wnt Signaling List” in Methods). Wnt signaling exhibited tissue- and directionality-specific adaptations in response to daily treadmill running (Figure 4A). Endurance training strongly upregulated the S_sec_ of positive Wnt regulators in scWAT, particularly from BAT, heart, and adrenal gland (Figure 4A). While training only modestly increased hypothalamic-targeting Wnt signaling from most origin tissues in general, hypothalamic-originating Wnt factors displayed a different trend, with negative Wnt regulators being more prominent in hypothalamus-to-peripheral circuits (Figure 4A). Assessing individual Wnt factors, *Rspo1* (R-Spondin 1; extracellular score: 4.7), a positive Wnt regulator, showed higher S_sec_ in most target tissues post-training (Figure 4B). The top 10 origin-target pairs with the greatest S_sec_ rank differences for *Rspo1* were all upregulated in TR8W, with spleen-to-key metabolic tissues - liver, adrenal gland, kidney, and scWAT - among the most prominent connections (Figure 4C). Wnt3a exhibited a distinct pattern, with its Ssec increasing across most target tissues following training, and vena cava emerging as a major origin tissue at TR8W (Figure 4B). Nine of the top 10 *Wnt3a* connections originated from vena cava, suggesting the circulatory system as a key *Wnt3a* source in response to endurance training (Figure 4C). Conversely, negative Wnt regulators such as Nog (Noggin) and Mdk exhibited reduced S_sec_ ranks in TR8W, suggesting a weaker endocrine role post-training (Figure 4C). Our findings suggest that Wnt signaling is associated with endurance training-induced endocrine remodeling, including metabolic adaptations in peripheral organs (e.g., skeletal muscle; Figure 3C) and neuroprotective mechanisms in the brain (Figure 3D-F). Positive Wnt signaling has been implicated in synaptic plasticity and neuroprotection, with its upregulation linked to protection against neuroinflammation and cognitive decline (30, 31). While these Wnt factors are confirmed to be extracellular and likely secreted, whether they cross the blood-brain barrier remains unknown. However, Wnt proteins can be secreted via exosomes (26), providing a potential mechanism for peripheral-to-brain signaling.

**Figure 4.**
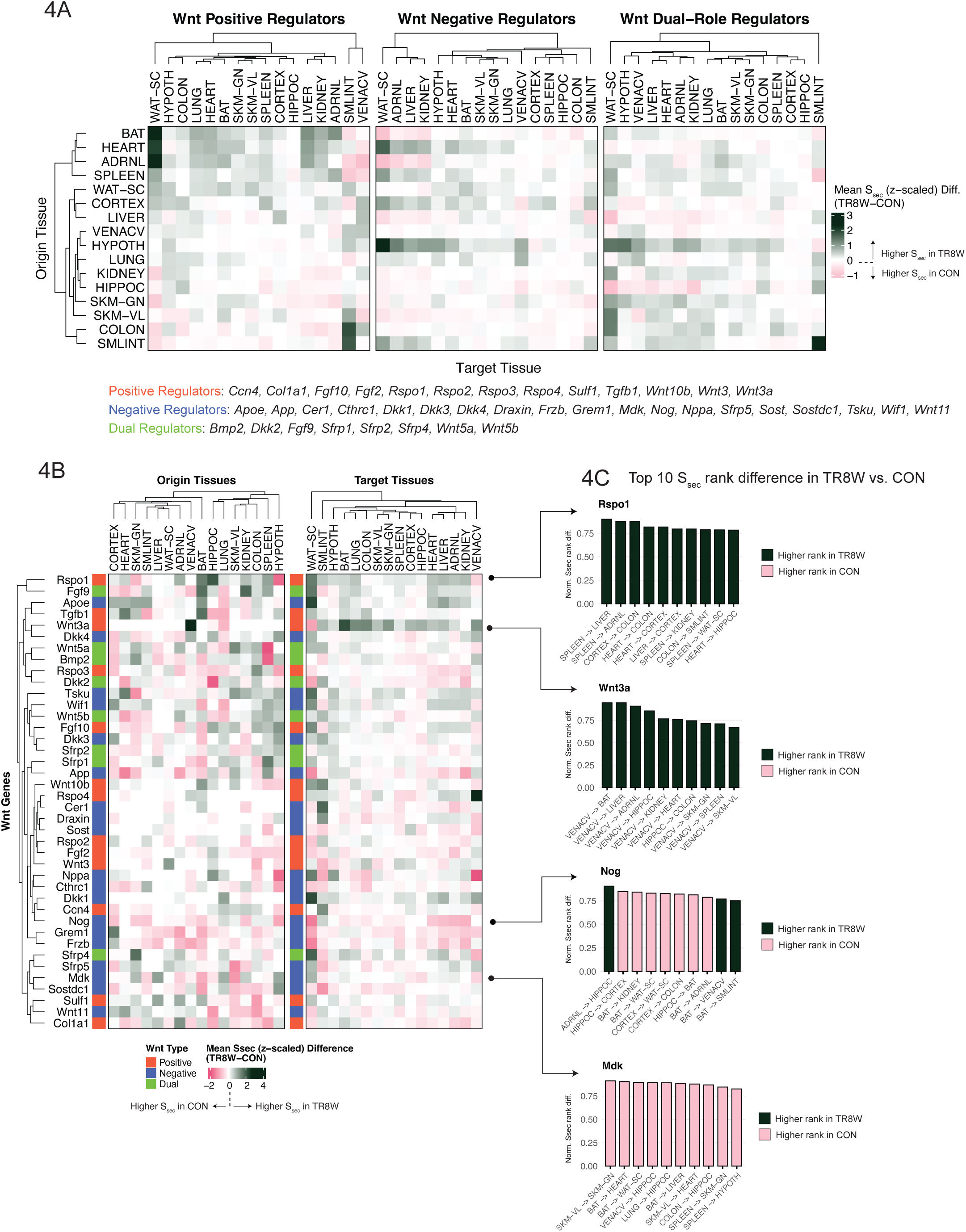
Feature- and tissue-specific remodeling of Wnt network by exercise training. A) Heatmaps displaying the average difference in S_sec_ (TR8W – CON) for positive, negative, and dual regulators of Wnt signaling. Each Ssec difference was z-scaled before calculating the average. Rows represent origin tissues, and columns represent target tissues. Member genes of the three categories are shown. B) Heatmaps illustrating the average difference in Ssec (TR8W – CON) for secretory Wnt factors, summarized by origin and target tissues. Each Ssec difference was z-scaled before averaging across origin or target tissues. C) Bar graphs showing the normalized S_sec_ rank difference between TR8W and CON for the top 10 origin-target tissue pairs with the largest rank differences for *Rspo1, Wnt3a, Nog,* and *Mdk*. Normalized rank difference was calculated by dividing the absolute S_sec_ rank difference between TR8W and CON by the total number of secretory genes in the origin tissue. The normalized rank difference ranges from 0 to <1. Dark bars indicate a higher rank in TR8W, while pink bars indicate a higher rank in CON.

## DISCUSSION

In this study, we leveraged an extensive dataset of transcriptomics and proteomics in 16 solid organs to interrogate inter-organ, gene-to-gene, and protein-to-protein correlations spanning more than 1,500 origin-target tissue pairs in a rigorously controlled preclinical endurance training model, Fisher 344 rat. Through the application of advanced systems genetics analytical approaches—QENIE (7, 8) and GD-CAT (11)—we quantified endurance training-induced adaptations in endocrine networks and inferred potential biological pathways that may be modulated by secretory features delivering, in part, health-related benefits of daily physical activity.

Our findings demonstrate that *in silico* estimations of crosstalk strength and endocrine responses align with well-established exercise-induced adaptations, such as leptin and IL-15 signaling, underscoring the validity of our approach. We provide evidence that endurance training induces extensive remodeling of endocrine networks, altering both overall inter-organ crosstalk strength and the individual contributions of specific secretory factors on tissue-selective pathways. Additionally, by meticulously filtering for known secretomes and inspecting their subcellular localization, we identified potential key secretory features that were most impacted by training in an origin-target-specific manner, particularly the involvement of Wnt signaling factors in brain physiology, offering a valuable resource for future mechanistic validation studies of novel and known exerkines. Importantly, this work expands our current understanding of the exerkine network by presenting a multi-tissue endocrine atlas that highlights potential mediators of exercise-induced tissue adaptations that are currently undiscovered.

Despite the unprecedented resolution of endocrine network adaptations provided by our study, several important limitations should be noted. As a correlation-based analysis, our approach cannot definitively establish causal relationships or exclude the possibility that observed S_sec_ changes reflect parallel exercise adaptations in both origin and target tissues, rather than direct endocrine signaling. Therefore, follow-up mechanistic studies are required to validate the functional roles of candidate secretory factors. While our well-controlled training model allowed for precise assessment of training effects on endocrine networks, the relative homogeneity of rats within each group may have limited biological diversity. For example, IL-6, one of the most studied exerkines, was only detected in a limited number of tissues (spleen and scWAT), restricting our ability to assess its systemic endocrine effects. Although the rigorous study design of MoTrPAC enabled profiling of the temporal dynamics of endocrine networks during training (i.e., 1 to 4 weeks), interpretation of these temporal changes should be made cautiously due to the absence of untrained control groups at intermediate time points.

Collectively, our integrated systems genetics approach using QENIE and GD-CAT revealed that endurance training drives widespread remodeling of inter-organ endocrine communication. By capturing both the strength and specificity of cross-tissue secretory relationships, we uncovered novel candidate endocrine factors and biological pathways that may underlie systemic adaptations to chronic exercise. These findings offer a foundational endocrine interaction map— linking molecular signals from origin to target tissues—that both confirms known biology and proposes new axes of crosstalk for mechanistic investigation. This resource provides a framework for future studies aiming to unravel how exercise-induced secreted factors orchestrate whole-body physiological adaptations.

## METHODS

### Molecular Transducers of Physical Activity Consortium (MoTrPAC) rat endurance training study

A detailed description of the endurance training, biochemical analyses, and differential analysis is available here (12). Briefly, six-month-old male and female Fischer 344 rats underwent progressive treadmill-based endurance training for 1, 2, 4, or 8 weeks (n=10 for each group; TR1W, TR2W, TR4W, TR8W). Tissues were collected 48 hours after the final exercise session. Age- and sex-matched sedentary rats served as untrained controls (n=10), whose tissue collection was synchronized with the 8-week training group. All normalized quality-controlled transcriptomics and proteomics datasets and differential analysis results were acquired using the MotrpacRatTraining6moData R package (https://github.com/MoTrPAC/MotrpacRatTraining6moData).

### Quantitative Endocrine Network Interaction Estimation (QENIE)

A detailed description of the original method is available here (7). Gene-to-gene and protein-to-protein biweight midcorrelation was performed on all possible tissue pairs per training condition. Resulting p-value matrices were filtered for known secreted proteins using the Uniprot rat secretome database (n=1321 unique features). Ssec was calculated as the sum of -log(p-value) for a given secretory feature in the origin tissue, correlating with all features in the target tissue. The sum was divided by the number of features in the target tissue for scaling.

### Gene Derived Correlation Across Tissues (GD-CAT)

From the biweight correlation between a given secretory feature and all target tissue features, target tissue features were ranked by the bicor value. Fast Gene Set Enrichment Analysis (fgsea R package (32)) was conducted on the ranked features to test for enriched biological pathways. TMsig R package was used for fgsea visualization. Benjamini-Hoechberg multiple correction was applied and terms with BH-adjusted p<0.05 were considered significant.

### Weighted Gene Co-expression Network Analysis (WGCNA)

#### Module construction and intercorrelation

WGCNA was used to identify distinct, non-overlapping gene clusters (“modules”) within each tissue transcriptomics matrix. Feature connectivity was calculated by summing correlation strengths with all other features across concatenated sample groups (nL=L50). A scale-free, signed network was constructed using the lowest soft-thresholding power β that achieved scale-free topology (R² >L0.85). Modules were defined using a dynamic tree-cutting algorithm with a merging threshold of 0.3, applied uniformly across all tissues. Module eigengenes (MEs), representing the first principal component of each module, were extracted and pairwise inter-module correlations were computed using biweight midcorrelation. P-values were adjusted for multiple testing using the Benjamini–Hochberg method.

#### Overrepresentation analysis (ORA)

For biological interpretation, ORA was performed on each module using the hypeR package, with Gene Ontology (GO) as the reference database. GO terms with FDRL<L0.05 were retained and grouped into general biological themes based on keyword matching:

- Translation (ribosom|translation|translational)
- Mitochondria / OxPhos (mitochond|respirat|oxidative)
- Immune (immune|inflamm|interferon|cytokine|antigen)
- Neuronal (synapse|axon|neuron|neurotransmitter|dendrite|neurogenesis)
- Cell Cycle (cell cycle|mitotic|division|proliferation)
- RNA Processing (rna|splic|processing|mrna)
- Lipid Metabolism (lipid|fatty|cholesterol|adipose|lipoprotein)
- Glucose Metabolism (glucose|glycolysis|gluconeogenesis)
- Development (development|morphogen|patterning|organ morphogenesis)
- Apoptosis (apoptosis|cell death|programmed cell)
- Autophagy / Degradation (autophagy|lysosome|degradation)
- Epigenetic Regulation (nucleus|chromatin|epigenetic|histone|methylation|acetylation)
- Signal Transduction (signal transduction|receptor|mapk|akt|signaling pathway)
- ECM / Adhesion (extracellular matrix|ecm|adhesion|integrin)
- Hormone Signaling (hormone|steroid|androgen|estrogen)

For each broad category, the GO term with the highest –log(FDR) was selected as representative. The similarity between modules was quantified using the Jaccard index, calculated as the ratio of overlapping significant GO terms to the total number of union terms between modules.

### Extracellular score

Rat-specific subcellular localization data were obtained from COMPARTMENTS (22), which provides four annotations for extracellular localization: “extracellular region,” “extracellular space,” “extracellular exosome,” and “extracellular vesicle.” Each annotation is assigned a score ranging from 0 to 5, with higher values indicating a greater probability of extracellular localization. For each secretory gene identified through the QENIE pipeline, the highest score among these four extracellular annotations was selected as the “Extracellular Score.”

### Curation of secretory Wnt signaling lists

Gene sets for “Positive regulation of Wnt signaling pathway” (GO:0030177) and “Negative regulation of Wnt signaling pathway” (GO:0030178) were obtained from the Gene Ontology database. Both sets were filtered to retain only genes identified in the QENIE pipeline with an extracellular score greater than 4. Genes that appeared in both lists were categorized as “dual regulators,” while exclusive positive and negative regulators were curated by removing these overlapping genes from their respective lists. This process resulted in 13 positive regulators, 19 negative regulators, and 8 dual regulators.

### Statistics

Paired T-test was used to compare the average S_sec_ between TR8W and CON for selected secretory features. Wilcoxon signed rank test was performed to test S_sec_ rank difference between TR8W and CON for a given origin-target tissue pair. Effect size r was calculated from rank sum test (r = Z/√N) where Z refers to the z-score from the Wilcoxon test and N is the total number of observations. Benjamini-Hoechberg multiple correction was applied. All statistical analyses were conducted through R.

## DATA AVAILABILITY

All normalized quality-controlled transcriptomics and proteomics data are available in public repository (https://motrpac-data.org/). Calculated S_sec_, WGCNA data, Uniprot secretome database, and COMPARTMENTS scores are hosted on GitHub (https://github.com/ahnchi/MoTrPAC-PASS1B-QENIE).

## CODE AVAILABILITY

Codes used in this manuscript to perform WGCNA, QENIE, and GD-CAT are hosted on GitHub (https://github.com/ahnchi/MoTrPAC-PASS1B-QENIE).

## ACKNOWLEDGEMENT

The MoTrPAC Study is supported by NIH grants U01AG055135, U01AR071158, U01AG055137, U01AR071130, U01AR071133.

## AUTHOR CONTRIBUTIONS

CA conceived the study and analyzed the data. All authors have participated in data interpretation. CA and LMS drafted the work. All authors have participated in revising the work and approved the final version of the manuscript.

---

**Supplementary Figure 1.**
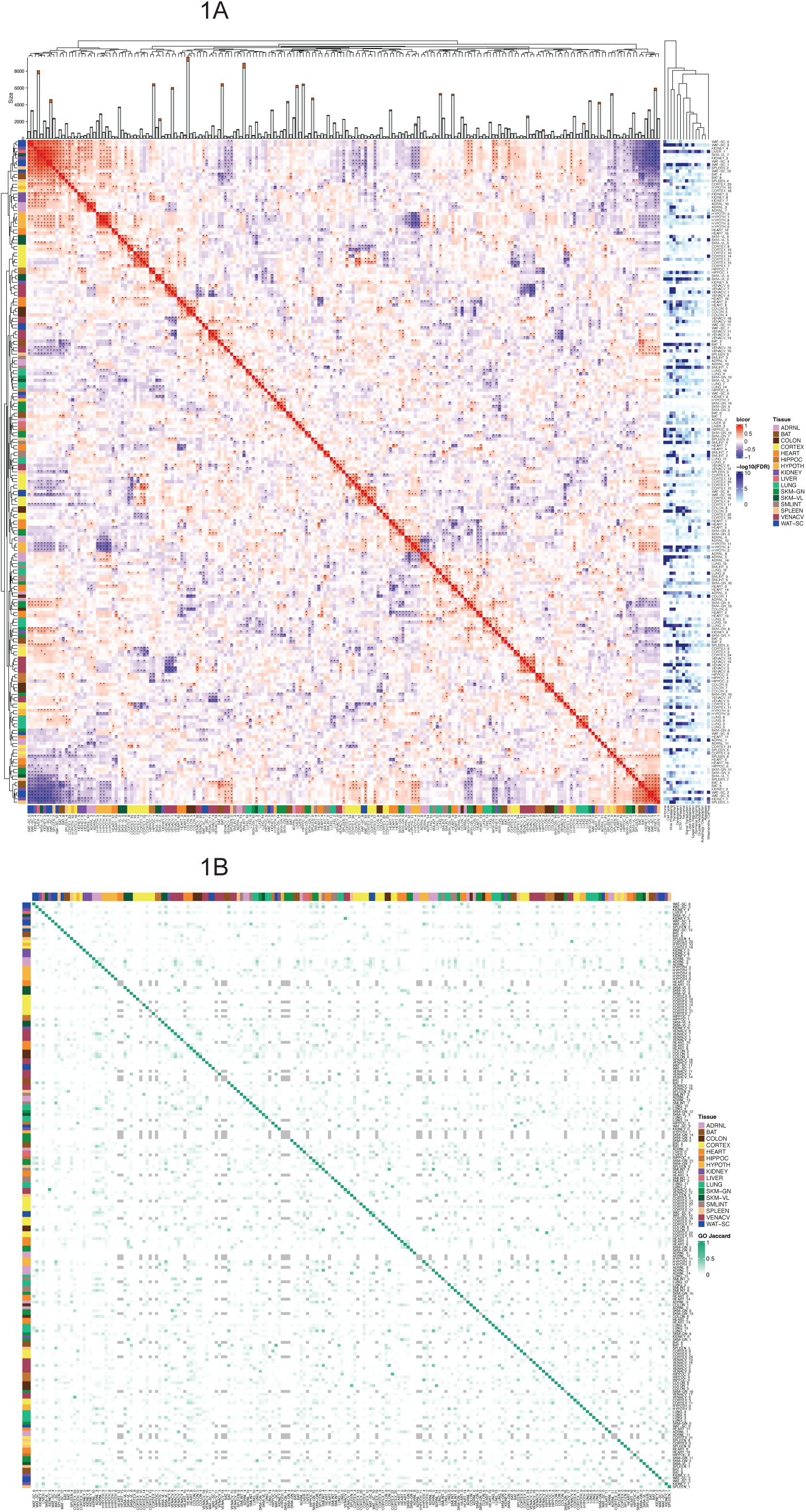
Module-to-module correlations across tissue transcriptomes. WGCNA was performed on concatenated gene expression matrices from all samples (n=50 rats) for each tissue, yielding a total of 203 co-expression modules across 16 tissues. A) ComplexHeatmap showing pairwise correlations between module eigengenes. *Adjusted p<0.05. Bar plots above each column represent module size (i.e., number of genes per module), with the orange portion indicating the number of secretory genes. Over-representation analysis (ORA) results are shown on the right. To reduce redundancy and highlight key biological themes, significantly enriched GO terms were manually curated into categories, including: Translation, Mitochondria/Oxphos, Immune, Neuronal, Cell Cycle, RNA Processing, Lipid Metabolism, Glucose Metabolism, Development, Apoptosis, Autophagy/Degradation, Epigenetic Regulation, Signal Transduction, ECM/Adhesion, and Hormone Signaling (see Methods for classification pipeline). B) Heatmap showing the Jaccard index (range: 0–1) for overlap of significantly enriched GO terms (FDR<0.05) between modules. Higher Jaccard values indicate greater overlap in functional enrichment. Row and column order mirrors that of panel A. Gray boxes indicate modules with no significantly enriched terms.

**Supplementary Figure 2.**
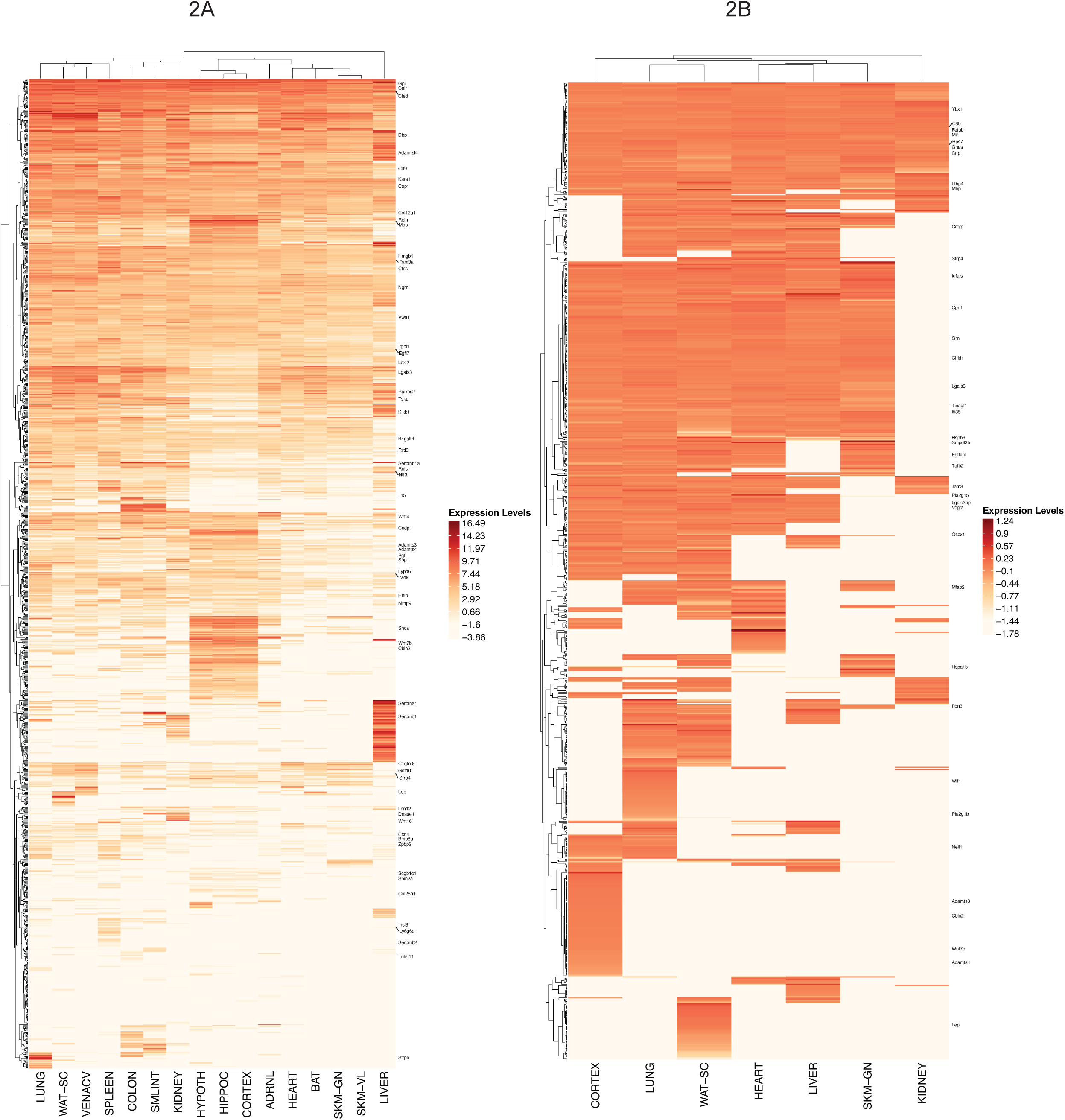
Secretory transcript expression and protein abundance across tissues. A) Heatmap of raw expression (log2 CPM) of 1117 secretory transcripts across 16 tissues. Undetected transcripts were imputed as the lowest expression level. B) Heatmap of raw abundance (log2 normalized abundance) of 797 secretory proteins across 7 tissues. Undetected proteins were imputed as the lowest abundance.

**Supplementary Figure 3.**
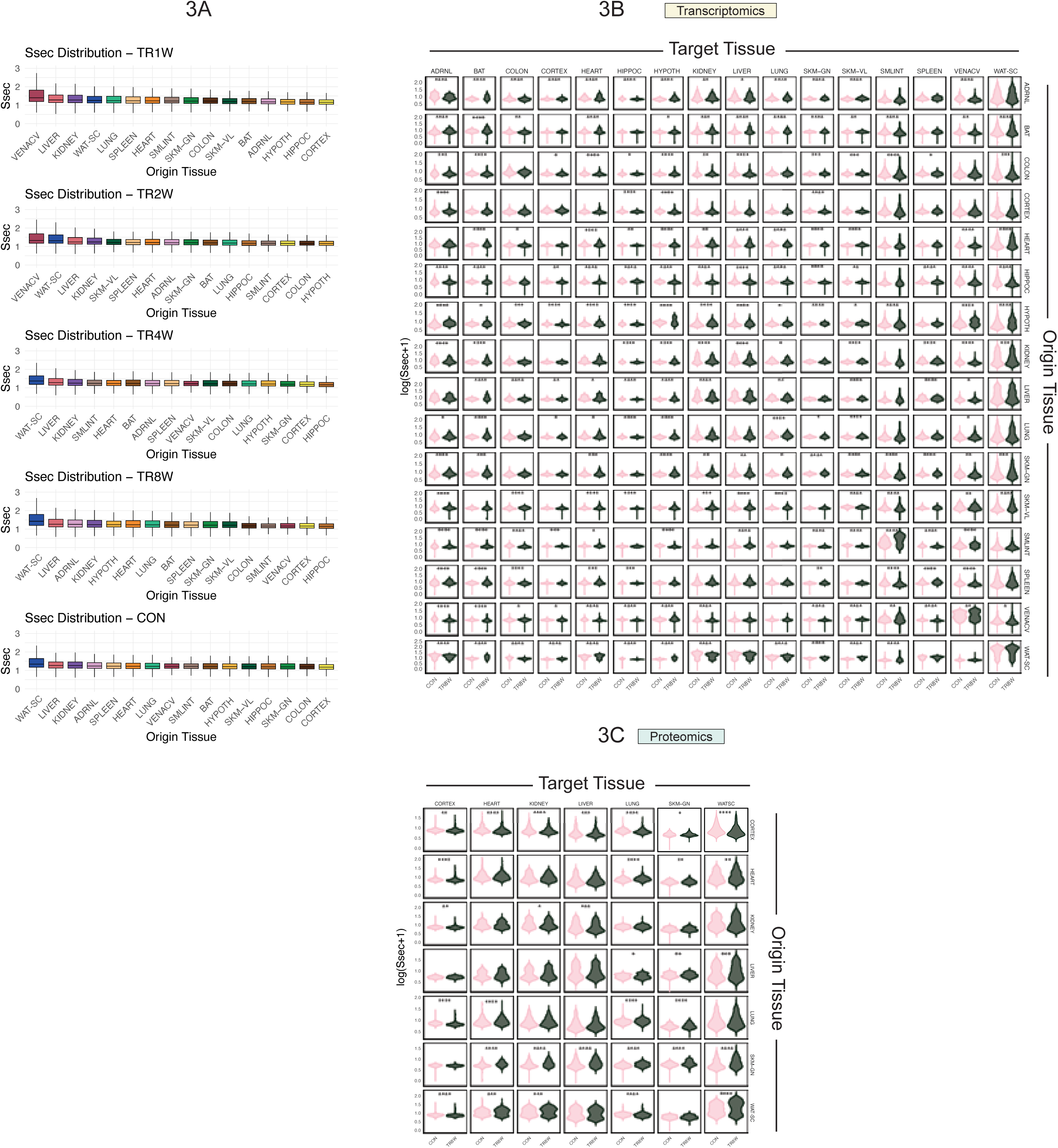
Ssec distribution over training and comparision of Ssec in CON vs. TR8W. A) Boxplots of tissue-specific Ssec across groups. B) Comparision of Ssec in CON vs. TR8W in all possible origin-target gene-to-gene correlation pairs. Paired T-test was performed on log(x+1) transformed Ssec. C) Comparision of Ssec in CON vs. TR8W in all possible origin-target protein-to-protein correlation pairs. Paired T-test was performed on log(x+1) transformed Ssec. *adjusted p<0.05, **adjusted p<0.01, ***adjusted p<0.001, ****adjusted p<0.0001. CON, control; TR1W, 1-week training, TR2W, 2-week training; TR4W, 4-week training; TR8W, 8-week training.

**Supplementary Figure 4.**
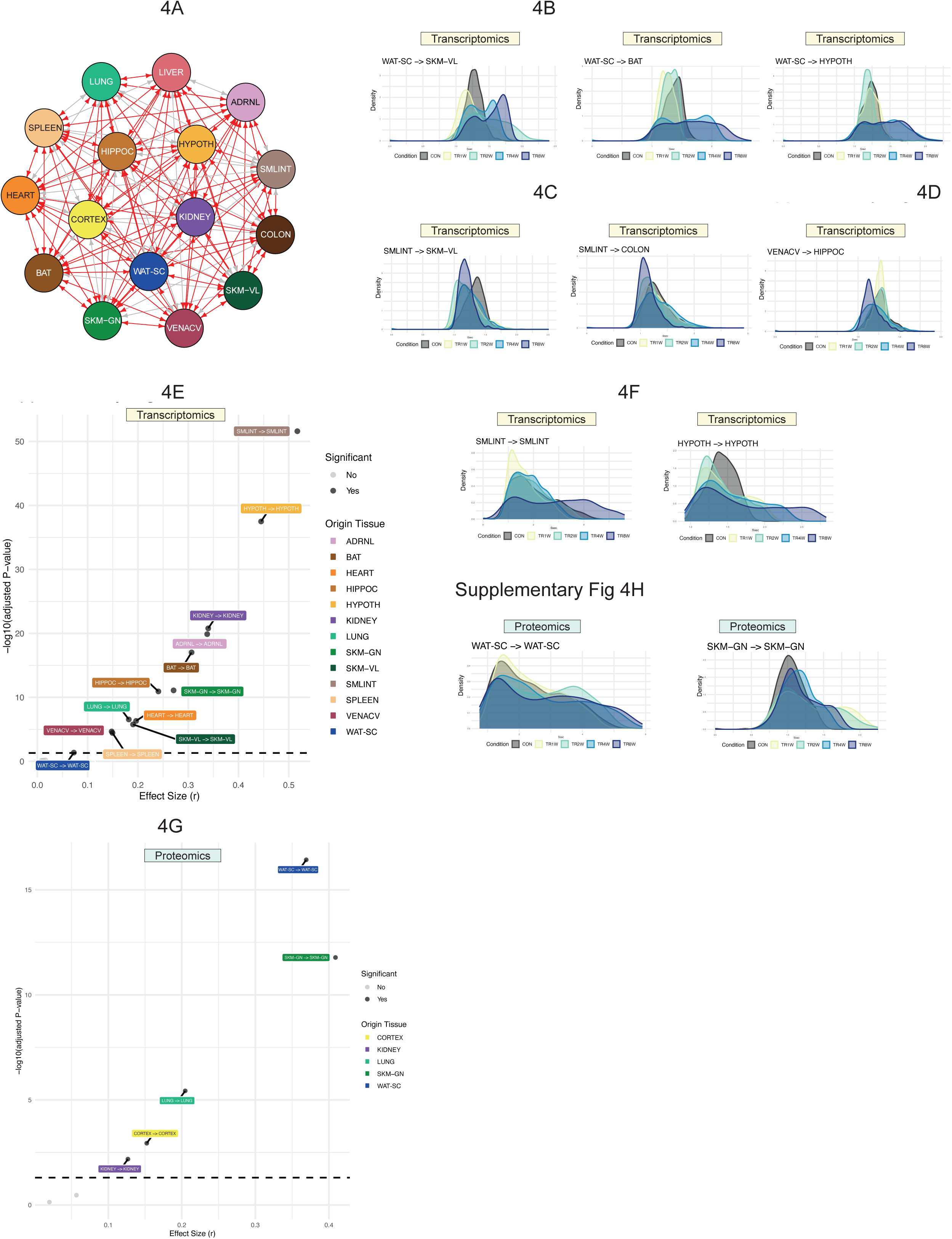
Comparision of Ssec rankings in CON vs. TR8W. A) Network plot showing significantly different Ssec ranking in origin-target pairs in CON vs. TR8W. Red arrows represent significant difference (adjusted p<0.05). B) Density plots of WAT-SC-originating (target: SKM-VL, BAT, and HYPOTH) Ssec across groups. C) Density plots of SMLINT-originating (target: SKM-VL and COLON) Ssec across groups. D) Density plots of a VENACV-to-HIPPOC Ssec across groups. E) Volcano plot of wilcoxon signed rank test results comparing gene-to-gene Ssec ranking of matched origin-target tissue pairs in CON vs. TR8W. F) Density plots of SMLINT-to-SMLINT and HYPOTH-to-HYPOTH Ssec across groups (gene-to-gene). G) Volcano plot of wilcoxon signed rank test results comparing protein-to-protein Ssec ranking of matched origin-target tissue pairs in CON vs. TR8W. H) Density plots of WAT-SC-to-WAT-SC and SKM-GN-to-SKM-GN Ssec across groups (protein-to-protein).

**Supplementary Figure 5.**
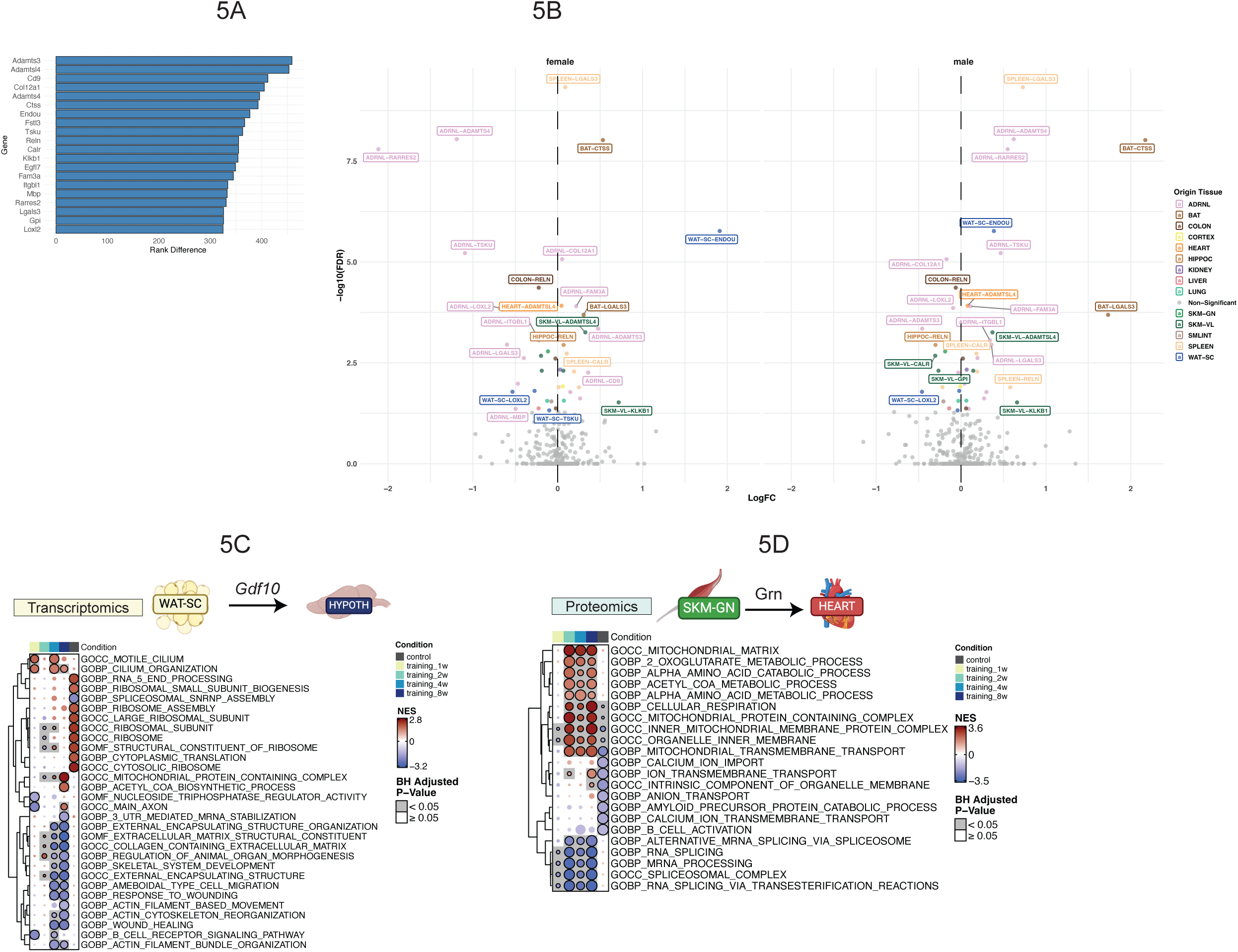
Top different global and tissue-specific secretory transcript. A) Top 20 global secretory transcripts with largest Ssec ranking difference between CON and TR8W. The ranking differences are shown in bar plots. B) Volcano plots showing differentially regulated transcripts (FDR<0.05) by 8-week training among top 20 global transcripts with most Ssec rank difference between TR8W and CON. Given that the differential analysis was conducted in each sex by MoTrPAC, volcano plots were created within each sex. Fgsea results of C) hypothalamus transcripts correlating with scWAT-derived Gdf10 and D) heart proteins correlating with gastrocnemius-dervied Grn.

**Supplementary Figure 6.**
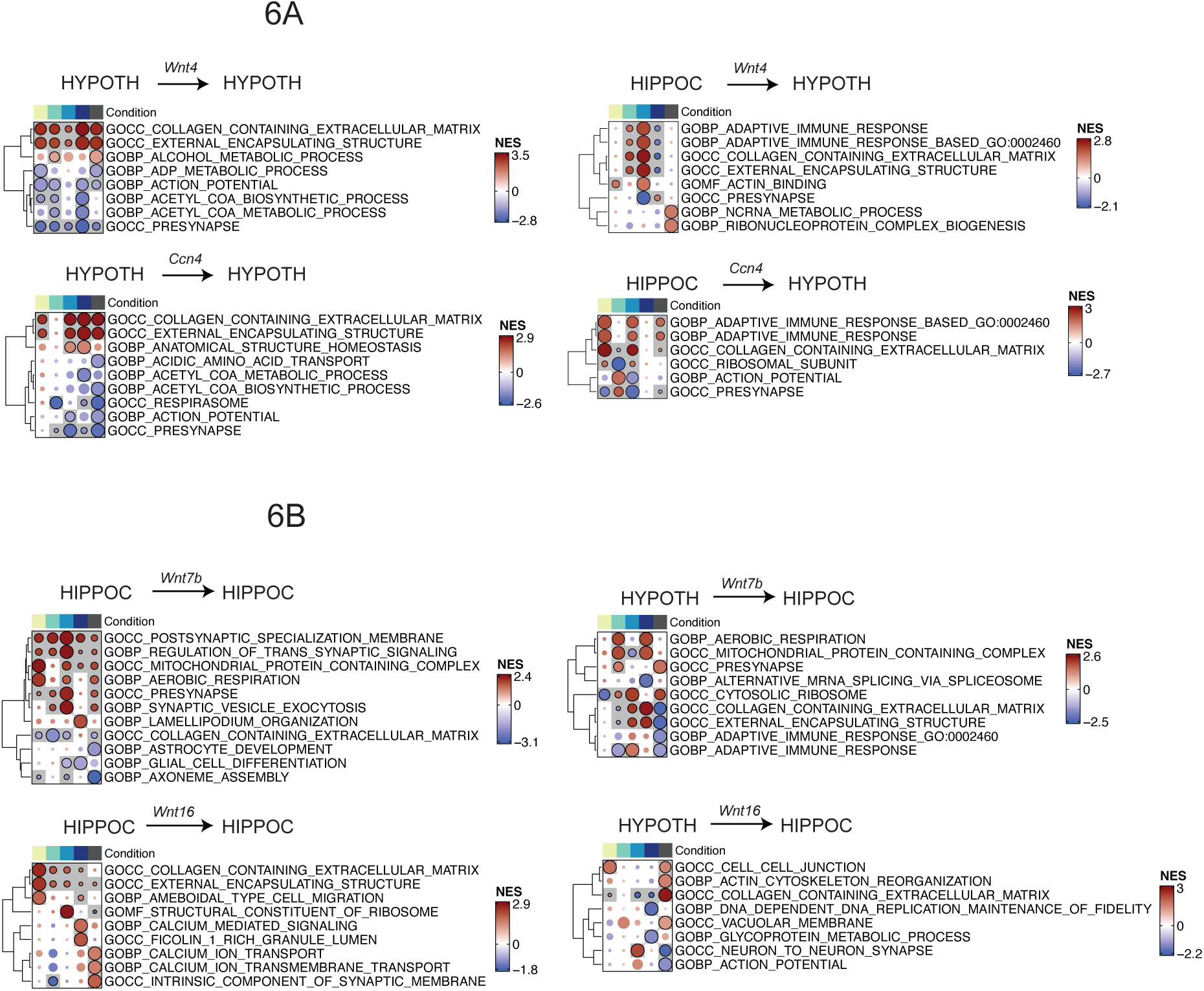
GD-CAT analysis of autocrine and paracrine signaling by secretory Wnt factors in the hypothalamus and hippocampus. A) Fgsea results of hypothalamus transcripts correlating with 1) hypothalamus Wnt4, 2) hippocampus Wnt4, 3) hypothalamus Ccn4, and 4) hippocampus Ccn4. B) Fgsea results of hippocampus transcripts correlating with 1) hippocampus Wnt7b, 2) hypothalamus Wnt7b, 3) hippocampus Wnt16, and 4) hypothalamus Wnt16.

